# The probability and duration of immigration in microbial communities

**DOI:** 10.1101/2024.12.12.628136

**Authors:** Thomas P Curtis, Ben Allen, Mathew Brown, Amy Bell, Donna Swan, Russel Davenport, William Sloan

## Abstract

Immigration is a fundamental feature of microbial communities. We propose a method to determine the probability (*P*_*i*_) *that* a number of immigrants (*i*) can attain an abundance *N* using the ratio of the probabilities of death *q* and division or “birth” *p* and the gambler’s ruin equation: *P*_*i*_*= (1-(q/p)*^*i*^*/(1-(q/p)*^*N*^). We estimate the probability of successful bioaugmentation or transplantation, the fate of a mutation infection and extinction. For example, an inoculum of 10^8^ bacteria with a *q/p* of 1.00000001 has a 10^-43^ chance of attaining an abundance of 10^10^. The immigration parameter used in neutral models, *m* is *1/(1-q/p)*. We calculated the long-term average value of m and *q/p* in a wastewater treatment plant. The value of *m* varies by >5 orders of magnitude, with a curious bimodal distribution. However, all the values of *q/p* are very close to, but greater than, *1*. We expect the long-term average value of *q/p* to be ∼1 in all stable microbial communities. In the absence of migration, bacterial populations with a *q/p* ≥1 will go extinct with probability 1. The link between *q/p* and infectious dose is known and we demonstrate that, in principle, the gambler’s ruin equation can estimate the infectious dose in naturally occurring infections, using *Vibrio cholerae* “carriers” to illustrate the point. We use the ratio *q/p* to estimate the time (measured in events or solar time) for a given change in abundance to happen. When *q/p=*1, extinction in even a small microbial population will take thousands of years.

## Introduction

In this paper, we wish to consider a simple model of the fate of a bacterium or bacteria of a given species when introduced into a microbial community. The kind of community we have in mind is a wastewater treatment plant or human gut. Microbial communities where the death of a microorganism is all but guaranteed and where “death” can be loss from the system or actual death by, for example, predation.

The arrival of individuals into a community is an unobservable but unavoidable feature of virtually all natural microbial communities.

For example, in engineering and medicine the deliberate addition of a species is referred to as bioaugmentation and probiotics, respectively. But there is, at present, no clear criterion for the success of this stratagem. In synthetic biology, the fear of the accidental immigration of a genetically manipulated organism into the environment has concerned researchers since the 1970s, but we have no way to evaluate the probability of such an event. In pathology, we know that successful immigration results in infection and, often, disease. The term immigration comes from ecology, and ecologists use the word invasion if the immigrant is novel.

These supposedly disparate phenomena are really all manifestations of the same process (Alexander 2023, Kinnunen et al 2016). A simple model of the fate of a bacterium or bacteria should offer insights into bioaugmentation, infection, and the fate of genetically manipulated microorganisms. Moreover, because local extinction is simply the reverse of invasion, we may also learn about eliminating a population.

However, such an understanding must be set in the context of contemporary ways of quantitatively describing the fate of an immigrant.

In ecology and microbial ecology, the probability of a death in a community being replaced by an immigrant is expressed as *m* (Hubbell 2001, Sloan et al 2006). Numbers are assumed to remain constant, so the probability of a death being replaced by a growth (called a birth) is, therefore, 1-*m*. The probability of success can be related to the number of immigrants with varying degrees of complexity (Blackburn et al 2015) (Leung et al 2004). The number of immigrants is sometimes called “propagule pressure” in ecological jargon.

The engineering community has other definitions, including “mass migration” (Frigon and Wells 2019), the proportion of the community in a reactor derived from the influent, and “net growth”, growth net of losses to predation and endogenous losses (not growth net of all losses) (Dottorini et al 2021) (Mei and Liu 2019).

In human, animal and plant pathology, infectivity is expressed as the 50% infectious dose (ID_50_), the number of microorganisms required to cause disease in half the exposed population. The ID_50_ can be thought of as a measure of immigration. ID_50_ values are determined experimentally from dose-response curves (Druet 1952, Feachem et al 1983). Quantitative studies of infection led Meynell to propose the “hypothesis of independent action” (Meynell and Stocker 1957, Meynell and Meynell 1970). The hypothesis that each invading bacterium has the potential to cause an infection. A number of researchers put this idea on a solid theoretical foundation in a suite of stochastic death/growth models (Shortley 1965, Saaty 1961, Williams 1965). However, these models proved difficult to calibrate because they used the probability birth and death independently (Shortley and Wilkins 1965). A number of authors noted that the ID_50_ could be inferred from the ratio of deaths to births, if the number of pathogens in in an infected subject were assumed to be very high (Shortley and Wilkins 1965, Williams and Meynell 1967). Subsequently, the hypothesis of independent action informed the use of dose-response curves in quantitative microbial risk assessment (Regli et al 1991) .

In this manuscript we (re)introduce the simple the gambler’s ruin equation. We then: i) Illustrate how the equation predicts the probability of immigration and extinction in two simple scenarios, Iii) Show how to estimate the key parameter, the ratio of death to birth, in two contexts in nature and (iii) use that ratio to estimate the time required for immigration or extinction to occur.

## Methods

The methods for collecting and analysing influent and mixed liquor of a full-scale wastewater treatment plant in Northeast England have been reported previously (Brown et al 2019). We used a longer time series (five years from June 2011 to June 2016 (258 sampling events)) in the same wastewater treatment plant and slightly refined bioinformatics. The methods are described in full in Appendix E.

## Results

### The Model

If we observe a microbe for long enough, sooner or later, it will do one of two things: reproduce with a probability *p* or die with a probability *q*. The probability of doing neither is *1-p-q* and so entirely defined by the other two probabilities and for the purpose of this model, *1-p-q = 0*. We are interested in the probability that the descendants of that microbe ultimately attain an abundance of N or alternatively, go extinct. The chosen abundance N can be an arbitrary parameter, an engineering goal, or the carrying capacity (often represented by a K in ecological models (Maynard-Smith 1974)).

This is a classical scenario in probability and Markov chains, often known as “the gambler’s ruin problem”. So called because it is like entering a casino with *i* francs (Pascal is credited with solving the problem) and playing a game with probability *p* and *q* of winning and losing a franc, respectively. A solution can be found in Appendix A and classical texts (Ross 2003).

Let *p*_*i*_ be the probability that the population starting with *i* individuals will eventually reach N. If *q*/*p* is not equal to 1, that is, the probability of a single organism doubling is greater or smaller than the probability of it dying, then:

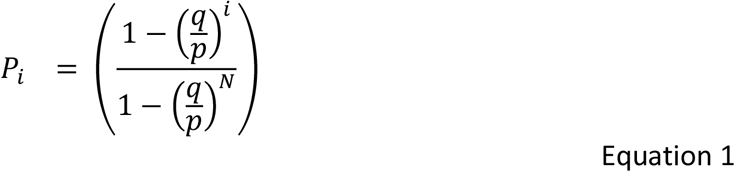

If p/q is = 1

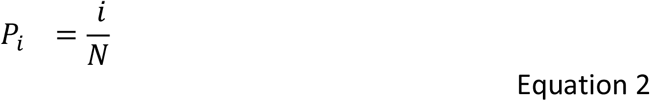

The model expresses the obvious simply. If birth is more likely than death, the probability of successfully immigrating is higher. If the ratio of deaths to births is greater than 1, successful immigration becomes quite unlikely.

An interesting corollary is that if *q/p* ≥1 and the gambler plays forever (the equivalent of setting N to infinity), the probability of extinction will become 1.

Superficially it would appear to be a simple, even trivial, binary; if q/p is less than 1 immigrations and if it is greater than 1 it does not. But for values close to 1 (which are, as we will show, the norm) a subtler picture emerges.

These subtleties become apparent when we examine the model in separate contexts: mutation, bioaugmentation, continuous flow systems infection. Finally we will use the ratio of deaths to births to consider the time for a change to take place.

### Mutation: where i=1

Consider the case of *i* is equal to 1. That is, there is just one immigrant. This is probably an extremely common event. Whilst it is unlikely that anyone would add a single bacterium into an environment, a mutant will inevitably arise with ∼ 1/300 births. Since most mutations will be unique, this is the equivalent of invasion by a single organism. In a system with 10^18^ individuals and a growth rate of 0.1 d^-1^ there will be ∼3 x 10^14^ such events a day. With so many events, “low probability” events can be of interest.

Thus, it is quite possible that a fraction of a slightly disadvantaged (∼*q*/*p* < 1.0001) bacteria generated each day by mutation can attain a modest abundance.

In larger (10^13^-10^18^ bacteria) communities, there may always be a significant background of mutations ready to be selected, should circumstances allow. The modest abundance they attain by chance could reduce the time it would take for them to become abundant and thus of functional importance in a system. Obviously, more disadvantaged mutations (*q*/*p* >1.05) are unlikely to attain even a modest abundance. For example, a single mutation with a *q*/*p* of 1.05 has less than a 10^-21^ chance of achieving an abundance of 10^3^. It would be helpful to know more than we do about the spectrum of fitness effects of mutations in bacteria.

### Probiotics and Bioaugmentation: where i>1

In medicine and environmental engineering, we may wish to add bacteria to a community in the hope that they will attain sufficient abundance to impact the ecosystem positively. However, a practitioner proposing such a course of action will presumably wish to have a high (say 90%) chance of success. Success being becoming sufficiently abundant to have the desired impact on the system. Success is context specific. However, we will assume success requires forming 0.1% of the biomass. In a biological treatment plant with 10^18^ individuals, this would imply a value of N of 10^15^

It is essentially impossible for an organism with a *q*/*p* value of >1 to attain a meaningful abundance. For example, an inoculum of 10^8^ bacteria with a *q*/*p* of 1.00000001 has a 10^-43^ chance of attaining an abundance of 10^10^.

Where death and growth match perfectly, *q*/*p* is exactly 1. Which seems unlikely, we can use equation 2 to see that:

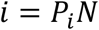

So, to have a 90% chance of success (*P*_*i*_=0.9), the inoculum (*i*) must be 90% of the desired final abundance (N). A proportion more akin to a transplantation than a bio-augmentation.

Where q/p is less than 1, success is not assured, but the dose required for a 90% chance of success can be estimated (Figure 2) by rearranging equation 3 and setting P_i_ to the desired probability.

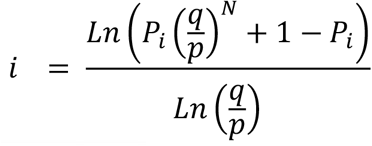

**Figure 1.**
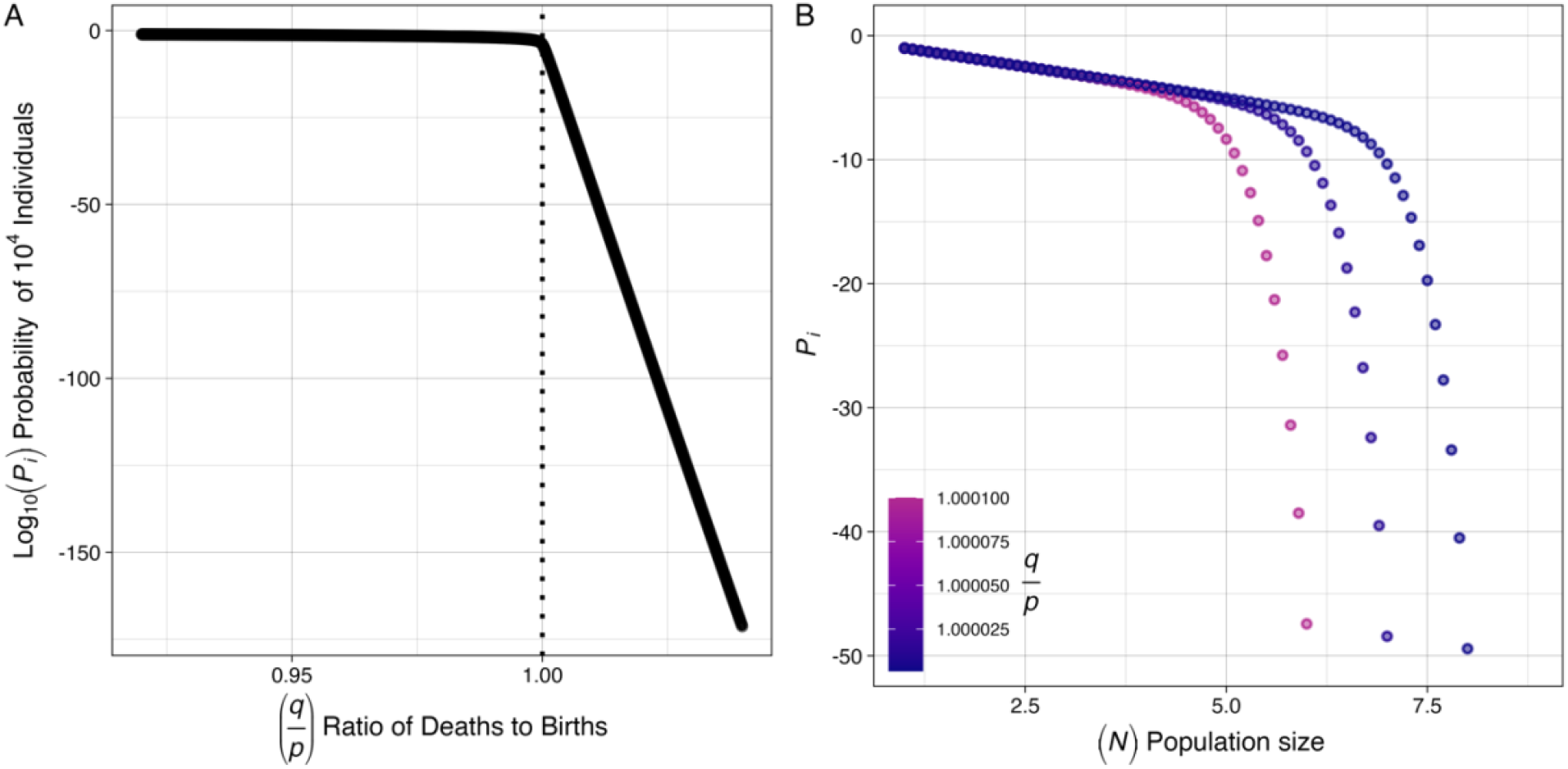
*A The probability of a single bacterium (i=1) achieving an abundance of 10*^*4*^ *(N= 10*^*4*^*) for a plausible range of ratios of death to growth B)* The log_10_ probability (*P*_*i*_) of a single immigrant or mutant (*i*=1) with a slight disadvantage (*q*/*p* =0.0001) attaining an abundance *Ni*.

**Figure 2.**
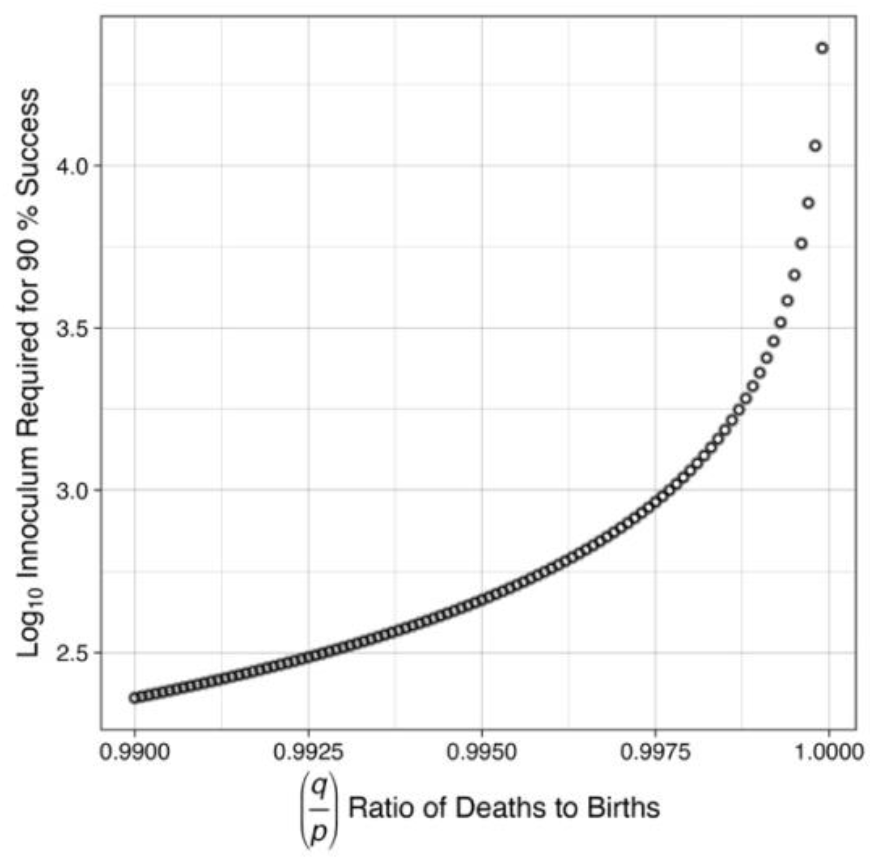
The inoculum required for a 90% chance of success for a given ratio (q/p) of deaths to births, assuming the desired abundance, N, is 10^6^.

A practitioner can increase their chances of success and reduce the inoculum required by reducing the ratio of deaths to births (Figure 2). The desired ratio or inoculum can now be calculated.

### Estimating the value of q/p in an engineered biological systems and microbial ecology more generally

In microbial ecology, immigration is expressed as *m*, the probability of a death being replaced by an immigrant. Typically, the value of *m* has been inferred by fitting a model to data (Harris et al 2017, Sloan et al 2006) as the compound parameter *N*_*T*_*m*, where *N*_*T*_ is the total number of bacteria in the system. Recently, Sloan et al. (Sloan et al 2021) demonstrated that *m* could be calculated directly in a chemostat. We have extended this reasoning to the aeration basin of a full-scale wastewater treatment plant and, in so doing, show a simple relationship between *m* and the parameter 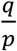.

The change in the number of a particular type of bacteria in a system is simply the sum of the number entering, the number dying (or leaving) and the number growing.

If *X* is the abundance of a particular bacterial species (cells/m^3^) (X_inf_ and X_reactor_, the concentration in the influent and mixed liquor, respectively), Q is the flow (m^3^/day) of influent entering the reactor, *μ* is the bacteria’s growth rate (d^-1^), *V* is the volume (m^3^) of the reactor and *θ* is the fraction of bacteria dying per day due to sludge wastage and other causes (d^-1^), then:

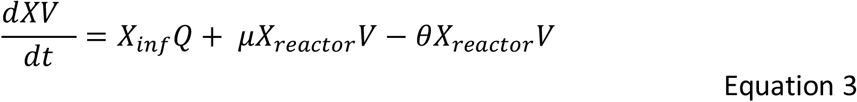

Equation 3 At steady state, and noting that hydraulic retention time (HRT) is V/Q, we can rearrange (see Appendix 2) equation 5 to express the parameters per death to show that:

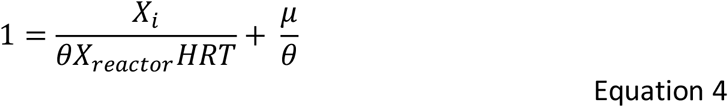

The parameter *m* is the number of immigrations per death, so at steady state:

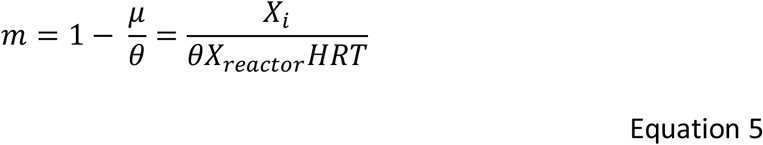

This equation shows the relationship between immigration, growth, death of biomass in the reactor, and the effluent and reactor conditions.

Thus, we have an equilibrium relationship between an equilibrium value for the migration parameter *m*, the ratio of births to deaths 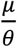, and therefore deaths to births 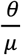, which gives an estimate of 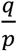:

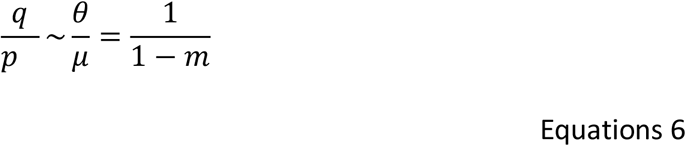

The value of *m* can be measured directly by measuring the influent and mixed liquor bacteria concentrations.

In a study described previously (Brown et al 2019) the number and diversity of bacteria in the influent and mixed liquor of an activated sludge plant were monitored weekly for 2 years. The mean sludge age in that period was 10 days, implying a death rate of 0.1, and the HRT was 12 hours. On this basis, we can calculate the values for *m* and thus 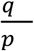.

By measuring the number of bacteria in the influent wastewater and the reactor weekly for 2 years and knowing the sludge age and HRT, we can estimate the value of *m* for many taxa. The range of values is large and bimodal (Figure 3, A). The taxa fall into two clear groups: those more abundant in the influent but less abundant in the reactor, and *vice versa* (Figure 4, A). However, any bacterium with an *m* value of <1 is growing in the reactor; thus, some taxa are growing unexpectedly, methanogens, (genus methanosphaera, m= 0.00008, genus methanobrevibacter, m= 0.0001) Some genera will be overlooked as detection limits will have set upper and lower limits on the observed values of *m*.

**Figure 3.**
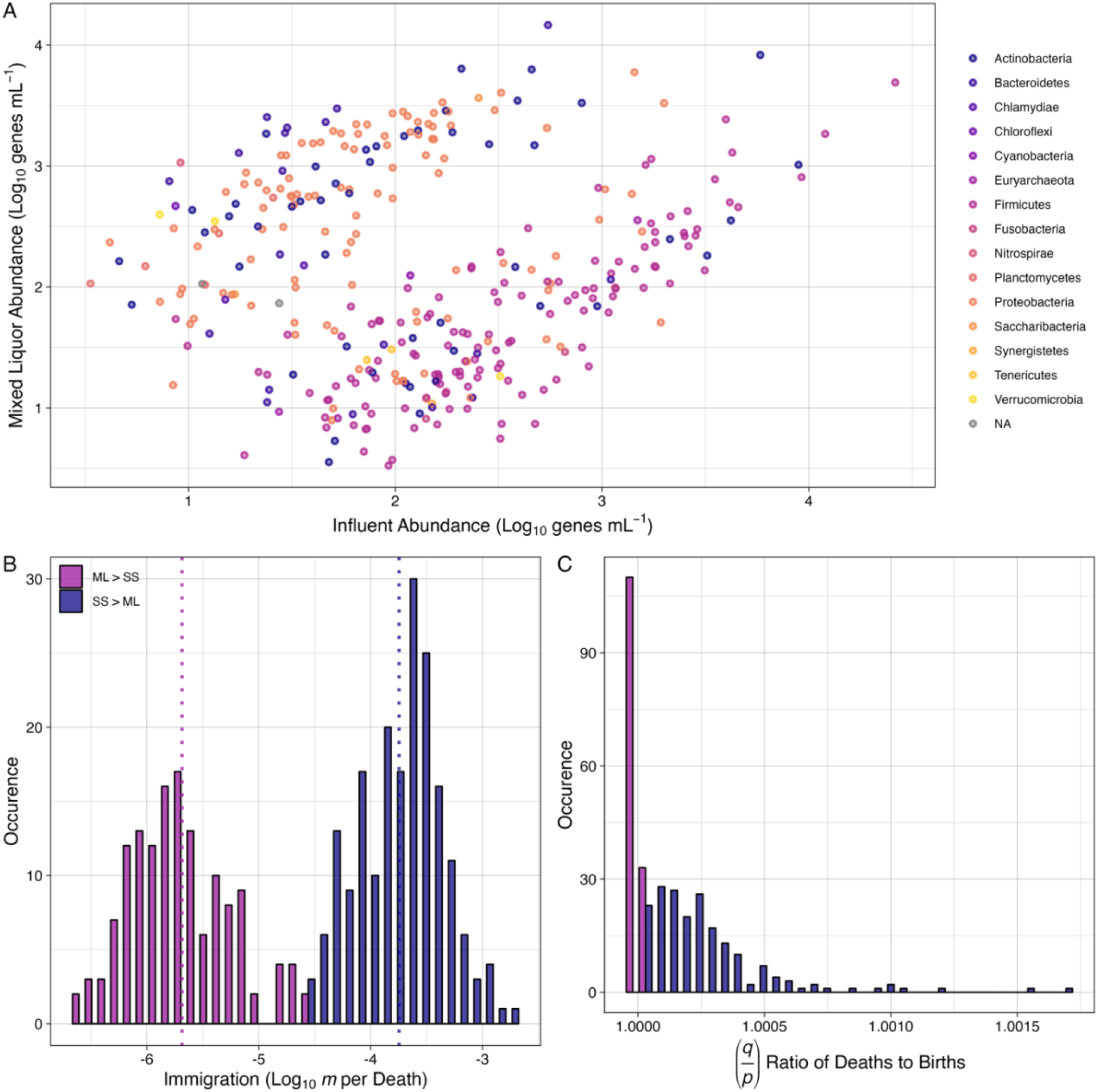
**A**. A comparison of taxa abundances in the influent and mixed liquor of the activated sludge plant. **B**. The bimodal distribution of immigration rates averaged over 2 years in an activated sludge plant: SS is settled sewage, ML is mixed liquor. **C** The distribution of values of the ratio in activated sludge.

**Figure 4:**
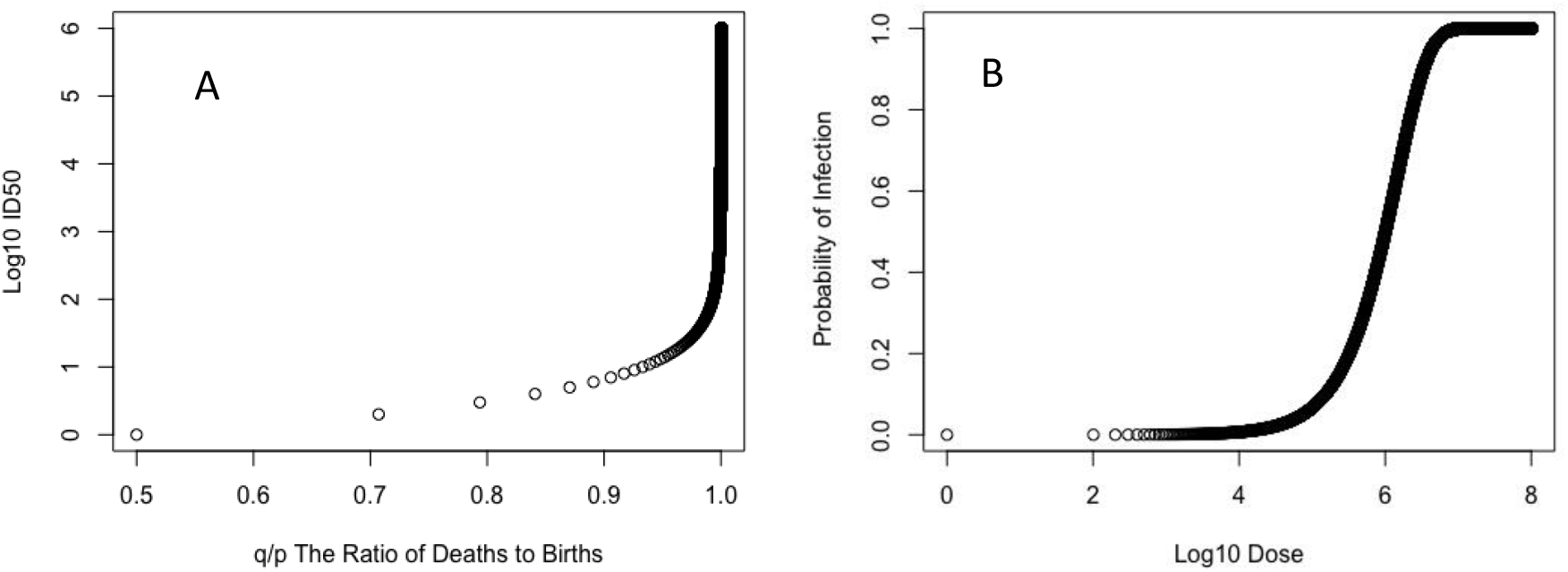
A) The relationships between the ratio of deaths to births and probability of infection (note actual values are less than 0.99999). B) A theoretical dose abundance curve for Salmonella, assuming a ratio of deaths to births of 0.99999931.

Nevertheless, using equation 5, we can see that the values of 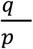 are very close to 1 (Figure 3C) and likely to range from 1.002 (from the genus Butyrivibrio_2) to 1.0000002 (from the genus Nitrospira)

### Estimating the value of q/p in Infections and estimating Infectivity

By contrast with the above a pathogen is axiomatically an organism with a q/p <1. (Meynell and Meynell 1970, Williams 1965). Moreover if a subject is made ill or killed by an infection the number of organisms infecting them will be very large., Where q/p is <1 and N>>1 the denominator in equation 1 quickly approximates to 1 and the Gamblers Ruin equation simplifies to the textbook relationships between the probability of infection and dose (Meynell and Meynell 1970)

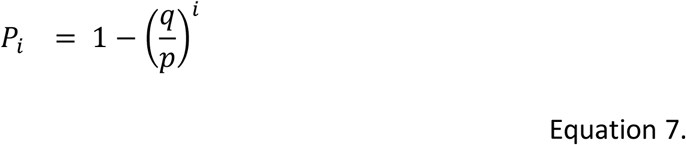

Thus, the infectious dose required to infect 50% of the subjects (ID_50_) of N, we can calculate *q*/*p*.

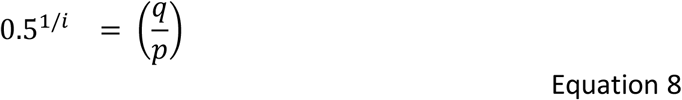

Which is an alternative to Shortley and Wilkins equation (q/p = 1-(0.69/ID_50_).

On this basis, one can calculate the relationship between *q*/*p* and ID_50_ (Figure 3 A) and conversely calculate the ratio of deaths to births from ID_50_. The ratio is close to, but less than, 1. Thus, in mice, Campylobacter serotype PEN 2 has an ID_50_ of 6.68 x 10^4^ (Blaser et al 1983) (Mitchell et al 2023), suggesting a *q*/*p* 0.9999896. In humans, Salmonellae have an ID_50_ of ∼10^6^ (Hornick et al 1966) (Feachem et al 1983), implying a *q*/*p* of 0.99999931. Whilst classical toxigenic *Vibrio cholerae* has an ID_50_ of ∼10^8^ (Feachem et al 1983), suggesting that *q*/*p* is 0.9999999931. Clearly, small changes to either deaths or births could be significant to infection.

Equation 4 (*P*= 1-(*q*/*p*)^*i*^) can be used to construct a plausible dose-response curve (Figure 4 B) and used in microbial risk assessment (usually as *P*= 1-e ^-*ki*^ where *k* is a fitted parameter)(Haas et al 2014). The fitted parameter *k* has a simple relationship with the theoretical parameter *q*/*p*:

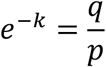

Estimating infectivity is problematic, especially in humans, as we usually rely on experiments on human subjects. Such experiments are ethically and scientifically challenging.

The Gambler’s Ruin approach might allow us to estimate ID_50_ and *q*/*p* by surveying populations, especially in diseases with both apparent and inapparent infections. Suppose that inapparent infections occur in people with ≤ N bacteria. The probability of being infected with at least N bacteria is:

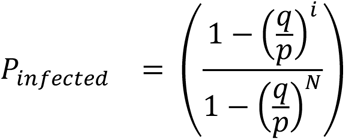

If we assume that to be symptomatic, you must have >>N bacteria, the probability of having symptomatic disease is:

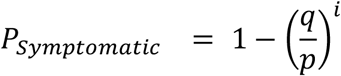

The fraction of symptomatic infections (*P*_symptomatic_/*P*_infected_) is:

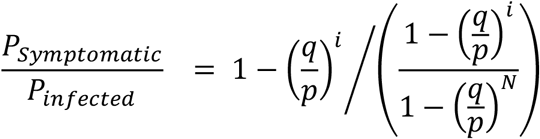

If we assume that both the symptomatic and the asymptomatic have received the same dose then:

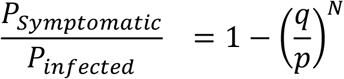

The ratio of *P*_symptomatic_/*P*_infected_ and the value of N can be determined by surveying a population. The value of *q*/*p* and thus ID_50_ (ID_50_ = ln (0.5)/ln (*q*/*p*)) can, in principle, be determined in a population by conducting quantitative surveys and without knowing the dose.

Dizon and colleagues (Dizon et al 1967) established that Cholera carriers (people with asymptomatic Cholera infections) had a geometric mean of 8.53 x10^3^ vibrios/gram in their stools. Assuming the mass of the contents of the large and small intestine is 1200g (Sender et al 2016), this gives a crude estimate of N of 7.3 x 10^7^. The ratio of infected-to-symptomatic ratio is purported to be between 1 and 30% (King et al 2008). This implies a range of *q*/*p* values from 0.999999936915 to 0.99999998351 and corresponding range of ID_50_ values from 1.1 x 10^7^ to 4.4 x10^7^ This is a tentative and preliminary “back of the envelope” calculation. However, the community ID_50_ estimates are between 10% and 40% of the values inferred from healthy prisoner experiments (Feachem et al 1983). This could imply that “real world” infectivity is higher than in healthy experiments or that those infected have a slightly higher dose than “carriers” or, as proposed (King et al 2008), the infected-to-symptomatic ratio is under estimated.

A pathogen can be eliminated by increasing the ratio of *q*/*p* to >> 1 (to ensure timely extinction) through a therapeutic measure, a physiological reaction or both. These values might be used to propose a therapeutic target or estimate the change in growth or death rate required to prevent or eliminate an infection. Clearly, to prevent infection, or at least make it less likely, one need only decrease the value of *i* through public health measures.

The same approach could be used to evaluate the safety margin for an organism, such as a genetically manipulated microorganism, that one might wish to go extinct. An organism with a *q*/*p* ratio of 1.1 is perhaps safer than one with a ratio of 1.00001.

### Time to attain a particular abundance

We now consider the time required for a particular change in numbers to occur. That is how long it would take to go from state 0 to state *n* or vice versa. Time in this context is typically measured in the number of deaths or births. This can be expressed as an expectation E[N_0,n_], the number of events it takes to go from 0 to *n* microbes.

When *q*/*p* is not equal to 1 Ross (Ross 2003) see Appendix, and other classical texts have shown that:

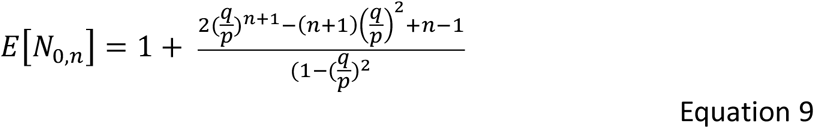

The expected number of events strongly depends on *q*/*p* (Figure 5), and at values <1 are linear with *n*. Though increases can, in principle, happen at *q*/*p* values > 1, they will take a long time.

**Figure 5.**
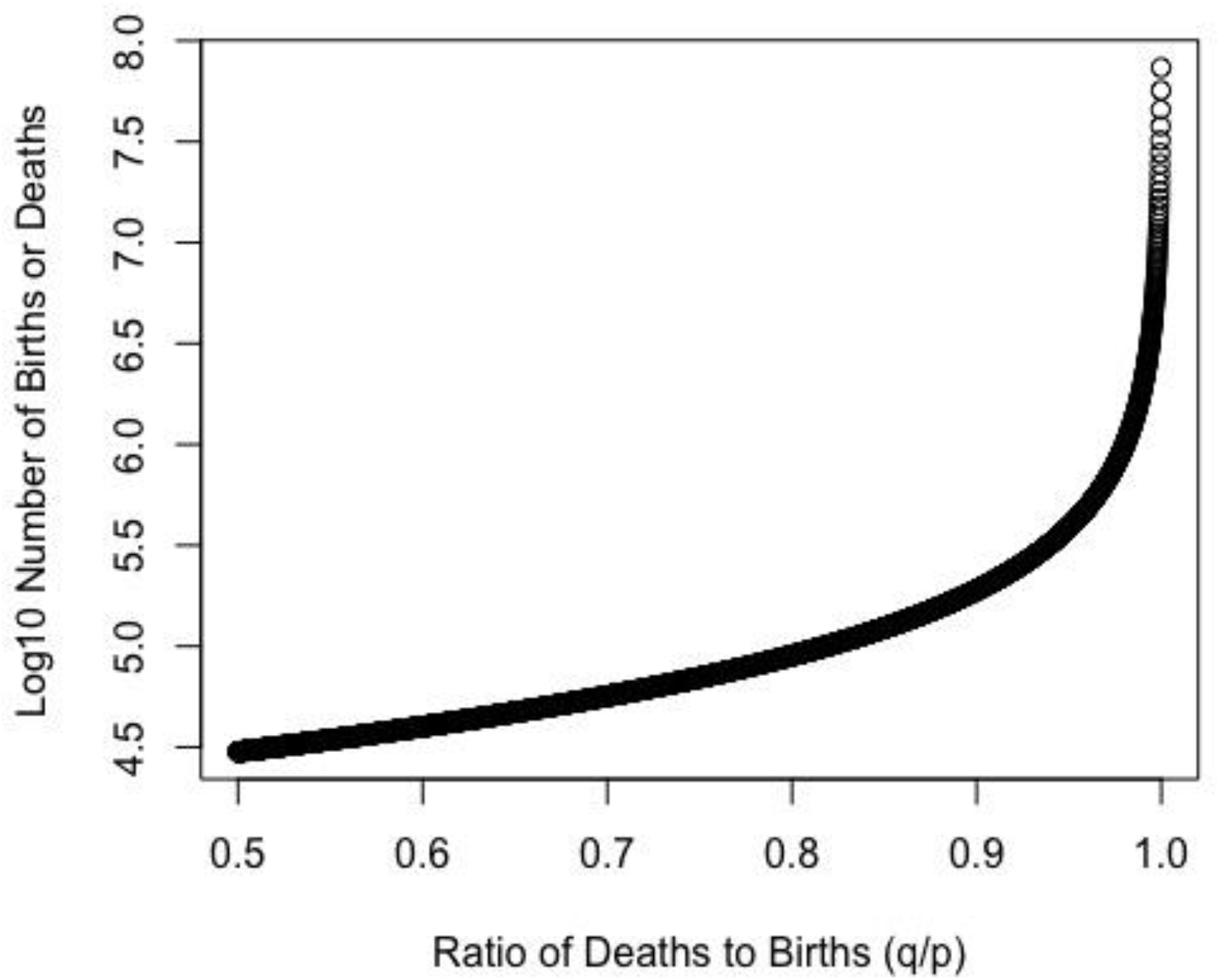
The Expectation of the number of events (births or deaths) that must occur for an organism to increase by 1000 individuals for differing values of deaths to births (*q*/*p*) of <1.

However, where *q*/*p* values are ∼1.

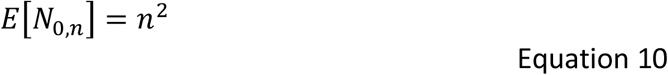

This is the expected number of events to go from a single immigrant to *n*, which is potentially very large indeed. Perhaps more importantly, as the number of events to extinction is:

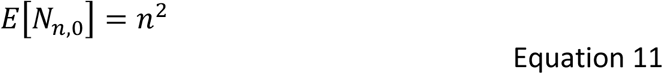

The standard deviation is also large (Ross proposes ∼0.8 n^2^).

This is interesting, as in stable ecosystems *q*/*p* is often ∼1. This could be a concern because the probability of extinction is 1 (from equation 1). How long will extinction take?

To answer this question, we must have a method to convert microbial time (events) to solar time (hours or days). The average time for 1 event (a birth or death) depends on the number of bacteria. If 10% of the population undergoes an event in a day and there are 1000 bacteria, then the mean time between events is 0.001 days, but if there are 10^14^ bacteria the mean time between events is 10^-13^ days. Thus, the mean time to an event will speed up or slow down as the microbial population increases or decreases.

To estimate that time, we have, in the first instance, adapted the approach described in Appendix 4 as follows.

Let M_i_ be the additional time that elapses from when a population enters the ith abundance class (i cells) until it has i +1 cells.

Then, M_0,n_ is the time required to get from state 0 cells to state n cells.

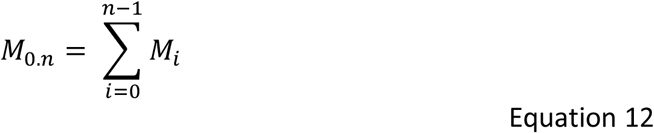

If *m*_*i*_ *= E[M*_*i*_] is the expectation of the time required to attain *i+1* after entering state *I*, we can show (appendix 5) that that expectation is:

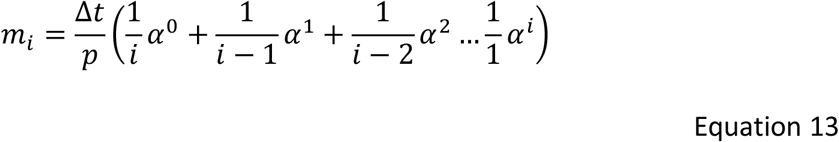

Where Δt is the time to doubling or death of an individual, and α is *q*/*p*. The effect of the ratio of *q*/*p* on time is very powerful (Figure 6).

**Figure 6.**
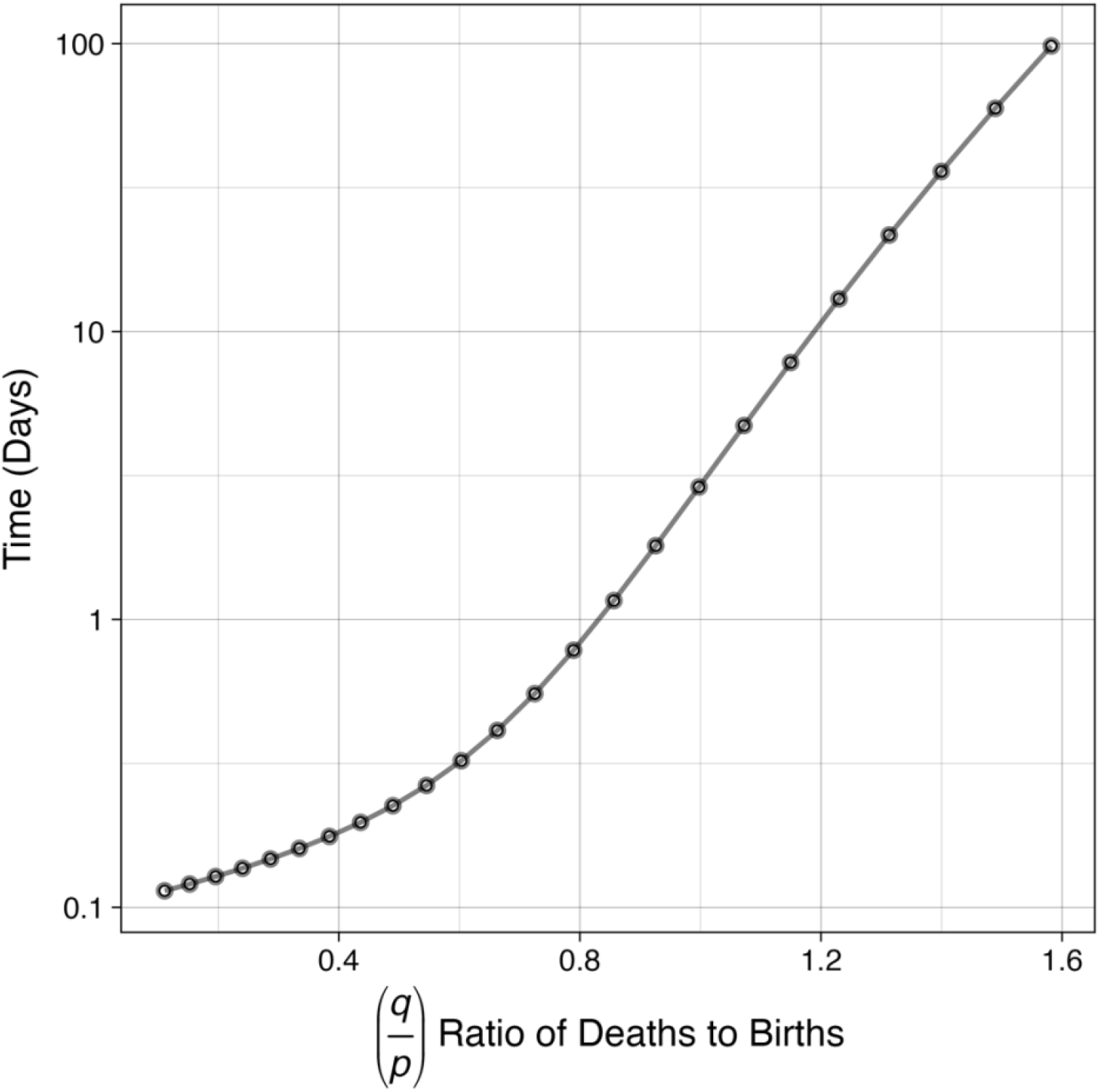
The expected time to increase by 1 bacterium when there are 10 individuals, and Δt is 0.1 days for differing ratios of deaths to births.

When α is 1, then equation 11 becomes a geometric series, and so:

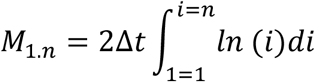

or

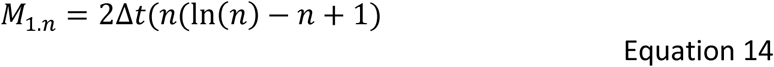

So, we can see that where *q*/*p* is 1, the change is slow. For a death rate of 0.1/day, it will take 10 million years to increase from 1 to 10^9^ (or vice versa).

Clearly, if change is to happen, α (q/p) must ≠ 1. We have not found an analytical solution for the sum of the terms in the series where α ≠1. However, where *ζ* = Δt/p, σ = α^-1^ it can be shown (appendix 4) that,

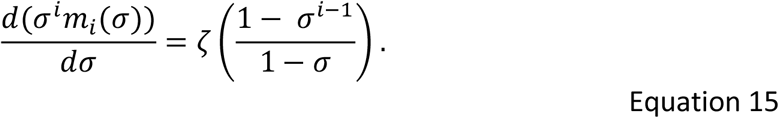

Now we can integrate numerically with respect to *σ* and divide by *σ*^*i*^ to get *m*_*i*_. It is then necessary to sum over all the values of m_i_ to obtain an estimate of time (Figure 8). Other methods can be found in the literature (Renshaw 1991).

**Figure 6.**
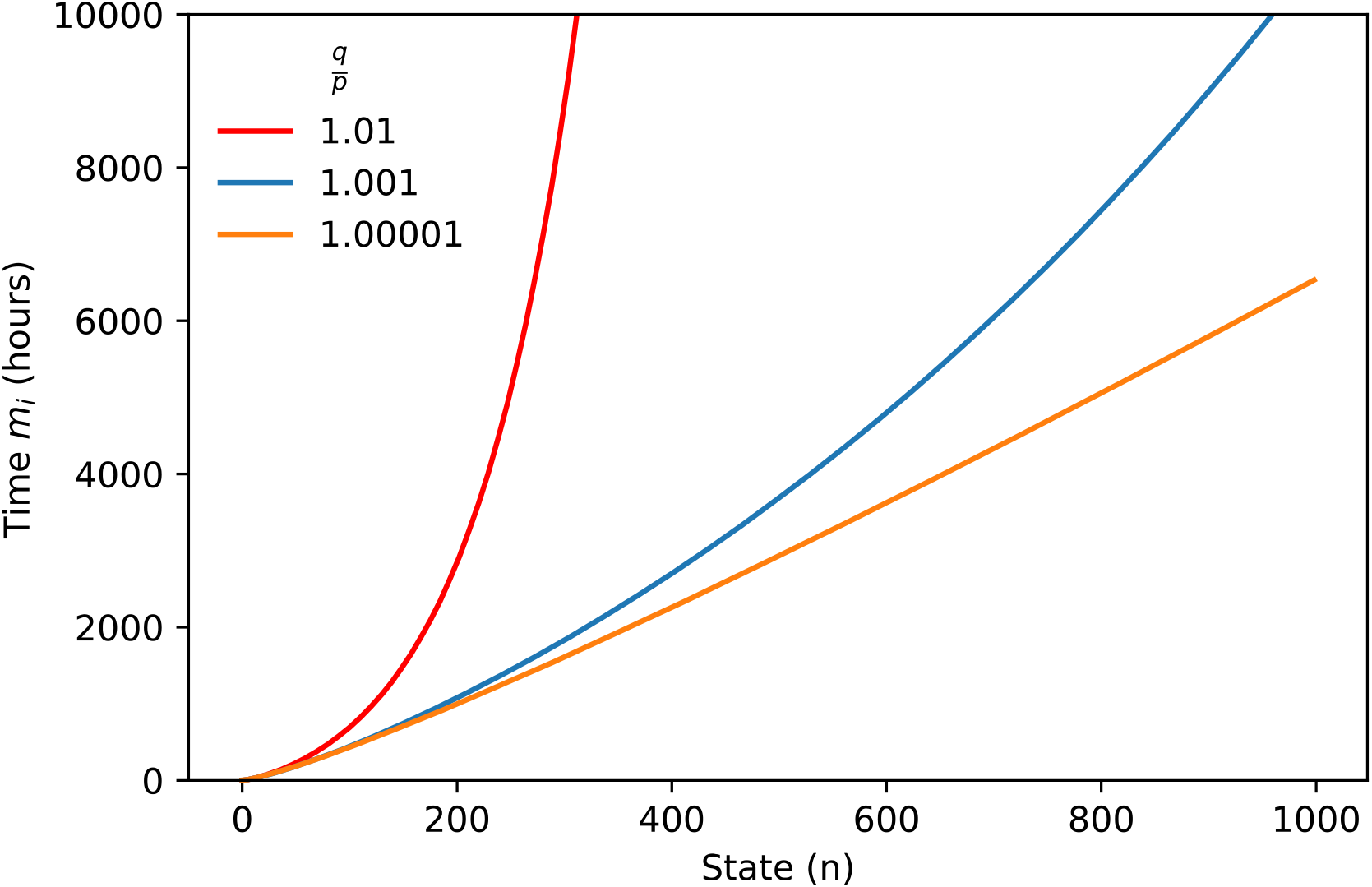
The time (hours) required to attain state N from a single organism with values of *q*/*p*>1 using equation 15.

## Discussion

We have revisited a simple and venerable mathematical relationship, the Gamblers Ruin equation and shown ways in which the key parameter, the ratio of deaths to births, can be evaluated in real microbial communities. The simple and parameterizable nature of this approach will hopefully mean that the relationships we propose can be applied to consider immigration, infection, bioaugmentation, extinction and other synonymous processes.

There are important *caveats* to this claim. Firstly, we have assumed that the ratio of births and deaths is constant. We defend this assumption as it is simple and, in many cases, will not affect the outcome or utility of the predictions. The second is that to predict the time for change when the ratio of births to deaths is not 1, we had to resort to a numerical solution which is not as elegant as we would like.

Stochastic models and the Gamblers Ruin equation are models not new. The pioneers in our field certainly (Saaty 1961), or almost certainly (Shortley and Wilkins 1965, Williams 1965) knew about this ancient expression. However, they did not explore its use. Perhaps because they were not aware of, or interested in, scenarios where probability of a given outcome was not basically 1. They were interested in the time from infection to disease (Williams and Meynell 1967). The early models required the user to determine the difference between the probabilities of death and growth; a number that is so close to zero it proved difficult to calculate from the animal experiments upon which they relied (Shortley and Wilkins 1965). The same reliance on the difference in deaths and births is found in subsequent stochastic models of microbial cultures (Coates et al 2018, Horowitz et al 2010). We are aware of an elegant study of the emergence of antimicrobial resistance in a petri dish (Saebelfeld et al 2022) does propose a synonymous equation, for the special case of an inoculum of 1, though ratio of deaths to births was not used or determined.

Other studies of invasion in microbial ecology eschew stochastic models in favour of qualitative assessments or multiple regression (Acosta et al 2015, Jones et al 2017, Kinnunen et al 2018). We argue that quantifying the basic mechanisms of invasion is preferable. Quantification permits a more nuanced, generalisable, and predictive understanding of the field.

For example, our finding that the ratio of deaths to births in stable populations in an established ecosystem (activated sludge) is ≥ 1. We admit that this is intuitively obvious and (given an assumption of constant numbers) arithmetically inevitable. However, it presents us with a paradox. In the absence of immigration, a stable population will go extinct ((Galton 1889, Kendall 1948, Lewontin and Cohen 1969, May 1971). Maynard-Smith (Maynard-Smith 1974) felt the inevitability of extinction to be a good reason to avoid stochastic models. We can resolve this paradox by invoking time. For even in the absence of immigration, extinction will take many thousands of years. Abundance offers some protection against extinction, locally, if not globally. Another way to avoid extinction is to simply not reproduce or die. For microbes time is measured in events, not solar years (George and O’Dwyer 2023); spore formation is simply a way to slow down time.

Stochastic models invoking immigration in microbial ecology are well established (Sloan et al 2006). We have made a link to such models (and reactor kinetics) by showing out the value of the immigration parameter m is linked to the ratio to deaths and births. Showing, amongst other things, that the value of m does not scale with reactor size; a finding we will explore in a subsequent manuscript.

The wide range values for m would seem to suggest that the notion of a dispersal and niche assembled communities (Loke and Chisholm 2023) could be discarded in favour of a more nuanced view of immigration as a continuum. However, the curious bimodal distribution of immigration parameters suggests exactly the opposite! Likewise, critics of neutral models can now point out that the value of *m* is a continuum that varies, by five orders of magnitude. The range of values of *m* may be greater, as some taxa were found in the reactor but not in the influent. However, supporters will point out that the ratio of deaths to births is almost always ∼1, supporting neutral theory’s assumption of equal fitness.

Moving away from theory, we would like to highlight some of the practical implications of our work.

Firstly, microbes can go extinct globally, and quickly, if the ratio of deaths to births is high. Global extinction might happen “by accident” if, for example, a chemical or class of chemicals, such as PFAS, were to poison a key enzyme such as ammonia monooxygenase (Curtis 2006). Where extinction is desirable, for example for an antimicrobial resistance gene or a genetically manipulated microorganism we can calculate the probability and duration of such an event

Stable communities are very difficult to invade, even by an organism that was perfectly adapted to that environment (q/p = 1), very large numbers are required. Thus the effects of bioaugmentation and probiotics will nearly always be transitory and will only be effective where the organism can have the desired effect before being made extinct. Often, it will be easier to change the environment to ensure *q*/*p* <1 than to design an organism with this property. Indeed, a change in environment seems to permit successful invasion in nature (Coyte et al 2021) and in engineering (Dybas et al 1998). The ratio of death to birth will be more likely to be <1 when a community is starting up. Consequently, seeding could be a better strategy for incorporating taxa into a community. We speculate that faecal transplantation often works because (amongst other things) relatively large numbers of immigrants are used, and the value of *q*/*p*, which in a stable community would be∼1, is reduced when the host is treated with antibiotics (Ghani et al 2022).

It may be possible to infer infectivity simply and ethically (where *q*/*p* is <1) by observing naturally infected populations. Our crude calculation of ID_50_ and *q*/*p* for *Vibrio cholerae* supports this proposal. However, we exploited an unusual disease and ancient datasets. Further work, more thought and better data and statistics are required before we can be more definitive about such an approach. More realistic dose-response curves might be helpful in quantitative microbial risk assessments. The assumption, that the dose is the same in both symptomatic and asymptomatic subjects is conservative. That is, it could lead us to overestimate the infectivity and so overstate the risk of infection, so even if it is wrong, it is safe.

Finally, we should not ignore the importance of low probability (eg 10 ^-14^) events. They will, for example, probably be fine tuning large wastewater treatment communities daily. Of course, the beneficial use of low probability events in engineering has already been proposed (Adams 1979) but not in the context of bacteria.

## Acknowledgements

We thank Northumbrian Water for their assistance and forbearance in gathering the wastewater treatment data. This work was supported by the UK Engineering and Physical Sciences Research Council EP/H012133/1 and EP/K039083/1 and the Biology and Biotechnology Research Council BB/Y512916/1. For the purpose of open access, the author has applied a Creative Commons Attribution (CC BY) licence to any Author Accepted Manuscript version arising from this submission. TPC would like to dedicate this manuscript to the memory of Professor Duncan Mara, public health engineer and classical scholar 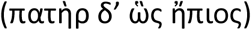, in thanks for his life and his guidance.

## Notes

### Competing Interest Statement

The authors have declared no competing interest.

