## Supplementary Material for "The probability and duration of immigration in microbial communities"

**Supplementary 1: Derivation of the Gamblers Ruin Problem (Ross 2003)**

In our idealised community a single bacterium will increase with the probability p ($\lambda$ in classical texts) and die with probability q ($\mu$ in classical texts) where q=1-p.

Let $P_i$ be the probability that an inoculum of size *i* will attain abundance *N* and *i* can be any number from 1 to *N-1*.

$$P_i = pP_{i+1} + qP_{i-1}$$

But *p+q= 1*

$$pP_i + qP_i = pP_{i+1} + qP_{i-1}$$

$$P_{i+1} - P_i = \frac{p}{q}(P_i - P_{i-1})$$

When i = 1, $P_{i-1}$ =0

$$P_2 - P_1 = \frac{p}{q}P_1 - P_0$$
$$P_2 - P_1 = \frac{p}{q}P_1$$

$$P_3 - P_2 = \frac{p}{q}(P_2 - P_1)$$

$$P_3 - P_2 = \frac{p}{q}(\frac{p}{q}P_1)$$

$$P_3 - P_2 = \left(\frac{p}{q}\right)^2 P_1$$

$$P_i - P_{i-1} = \left(\frac{p}{q}\right)^{i-1} P_1$$

Adding i-1 terms allow us to calculate:

$$P_i - P_1 = P_1\left[\left(\frac{p}{q}\right) + \left(\frac{p}{q}\right)^2 +..+ \left(\frac{p}{q}\right)^{i-1}\right]$$

The terms inside the square bracket are a geometric series so.

$$When \frac{q}{p} \neq 1 \ldots P_i = \frac{1 - \left({}^q/_p\right)^i}{1 - \left({}^q/_p\right)} P_1$$

Equation 1.1

$$When \frac{q}{p} = 1 \ldots . P_i = iP_1$$

Equation 1.2

$P_N$ is the probability of N bacteria attaining an abundance of N, which is obviously 1. Thus if $P_i = P_N = 1$ we can rearrange the equations above and calculate $P_1$ the probability of a single new bacterium attaining a density of N

If *p = q =0.5* then *q/ p = 1*

$$P_1 \quad = \frac{1}{N}$$

Equation 1.3

But if $q_\theta / p_\mu$ does not equal *1*

$$P_1 \quad = \left( \frac{1 - \left(\frac{q}{p}\right)}{1 - \left(\frac{q}{p}\right)^N} \right)$$

Equation 1.4

These two equations represent the probability of single new organism, for example a new mutation, acquiring an abundance of N.

We can determine $P_i$, the probability of i organisms attaining an abundance N is inoculated into by either:

Inserting Equation 1.3 into Equation 1.2 (were *q/p = 1*) to give:

$$P_i \quad = \frac{i}{N}$$

Or inserting Equation 1.4 into Equation 1.1 ($q_\theta / p_\mu$ *is not equal to 1*) to give:

$$P_i \quad = \left( \frac{1 - \left(\frac{q}{p}\right)^i}{1 - \left(\frac{q}{p}\right)^N} \right)$$

Thus the probability of an organism with a *q/p of less than 1* attaining an abundance of N is very low indeed even if N and i are very close.

Ross S. *Introduction to Probability Models*: Academic Press, 2003.
